# Releasable hydrogel microsphere-drug conjugates as generic prodrugs for long-acting intra-tumoral therapy

**DOI:** 10.1101/2024.02.16.580368

**Authors:** Jeff Henise, John A. Hangasky, Deborah Charych, Christopher W. Carreras, Gary W. Ashley, Daniel V. Santi

## Abstract

Intratumoral (IT) therapy is a powerful method of controlling tumor growth, but a major unsolved problem is the rapidity that injected drugs exit tumors, limiting on-target exposure and efficacy. We have developed a generic long acting IT delivery-system in which a drug is covalently tethered to hydrogel microspheres (MS) by a cleavable linker; upon injection the conjugate forms a depot that slowly releases the drug and “bathes” the tumor for long periods. We established technology to measure tissue pharmacokinetics and studied MSs attached to SN-38, a topoisomerase 1 inhibitor. When MS∼SN-38 was injected locally, tissues showed high levels of SN-38 with a long half-life of ∼1 week. IT MS∼SN-38 was ∼10-fold more efficacious as an anti-tumor agent than systemic SN-38. We also propose and provide an example that long-acting IT therapy might enable safe use of two drugs with overlapping toxicities. Here, long-acting IT MS∼SN-38 is delivered with concurrent systemic PARP inhibitor. The tumor is exposed to both drugs whereas other tissues are exposed only to the systemic drug; synergistic anti-tumor activity supported the validity of this approach. We propose use of this approach to increase efficacy and reduce toxicities of combinations of immune checkpoint inhibitors such as ***α***CTLA4 and ***α***PD-1.

## Introduction

Systemic administration of oncology agents can lead to severe adverse effects that limit the dose and hinder achievement of therapeutically active regimens. Tactics that address this problem often seek to localize the drug into or close to the tumor target with minimal systemic exposure. One very effective approach is locoregional delivery of the therapeutic agent by intra-tumoral (IT), peri-tumoral, or intra-lymph node injections (1-5)}. IT, or other locoregional injection of a drug provides very high local concentration – high efficacy – with a low systemic concentration – low toxicity. Although IT injections were once limited to accessible surface tumors, image-guided delivery has made the approach feasible across a range of target organs (6). Also, IT therapy has received much recent interest for immune-therapeutic and immuno-chemotherapeutic agents where local delivery could overcome the suppressive tumor microenvironment and activate antitumor immune responses (2). In spite of the potential advantages of IT therapy, in practice the rapid escape of most locally injected therapies undermines their potential advantages – for example, mAbs or ADCs with systemic t_1/2_s of several weeks have an IT t_1/2_ of only ∼6 hr (7, 8). We and others (e.g. Refs. (9, 10)) posit that the major reason IT therapies have not been more successful and accepted is their short IT residence time, and that long-acting IT therapies could play a transformative role in cancer therapy.

Several approaches have been used to prolong tumor residence time of IT-administered therapeutics. One is to simply increase the viscosity of the delivery system to retard diffusion from the injection site. For example, IT administration of a co-formulation of IL-12 with the viscous carrier chitosan increases the IT t_1/2_ of the cytokine from 3 to ∼15 hr, with marked improvement of tumor regression (11). Likewise, pluronic acid increased the IT t_1/2_ of a mAb from ∼6 to 20 hour which was accompanied by increased growth inhibition of tumors (7). But, tumor residence time of drugs using such agents is limited to ∼1 day because of dissipation of the viscous additive. Another approach to IT or intra-lymph node (5) half-life extension is to encapsulate the drug into ester-based polymeric microspheres (e.g. PGLA) that slowly release the drug upon hydrolytic erosion. While such carriers are quite effective for delivery of small molecules, the need for organic solvents during encapsulation, the acidic environment they create upon erosion, and chemical reactions with amino acid side chains make them unsuitable for use with proteins (12); indeed, cytokines lose significant activity during encapsulation in such polymers and upon storage (13, 14).

The most successful approaches for increasing IT retention of proteins involve some form of “anchoring” to local targets (10, 15). In one tactic, attachment of lipid groups to the drug (16) produced membrane-localizing forms with long-lasting effects. In another anchoring approach, IT injection of protein fusions containing components that bind extracellular matrix (ECM)-proteins such as collagen, fibronectin and others, lead to high IT retention. For example, appending a collagen-binding moiety to proteins anchors them to the tumor ECM and endows them with IT t_1/2_s of ∼24 hour (9). Likely the most advanced intra-lesional immunocytokines are those developed by Neri and Philogen (https://www.philogen.com) that anchor proteins to the extra domain B splice variant of fibronectin by the L19 antibody (17). IT administered L19-containing immunocytokines in late-stage clinical trials include Darleukin (L19IL2; NCT02076633) and Daromun (L19IL-2 + L19TNF NCT01253096). Finally, fusing an alum-binding moiety to therapeutic proteins provides a very effective way of anchoring proteins to the ECM by alum-tethering (18, 19)(https://ankyratx.com/). However, anchoring approaches generally require modification of native proteins and are not amenable for use with small molecules or peptides. Clearly, IT therapy would benefit from sustained delivery systems that are tumor agnostic, reliably provide prolonged and predictable drug release rates, and compatible with all classes of therapeutic agents – small molecules, nucleic acids, peptides and proteins.

We have developed a general approach for half-life extension of therapeutics that should be generally and directly applicable to IT drugs (20, 21). Here, a drug is covalently tethered to a long-lived carrier by a linker that slowly cleaves by β-elimination to release the native drug (**Scheme 1**). The cleavage rate of the linker is controlled by the nature of an electron-withdrawing “modulator” (Mod) which regulates the acidity of an adjacent carbon-hydrogen bond. These linkers can control drug release rates over long periods, are not affected by enzymes and are extraordinarily stable when stored at low pH and temperature (20, 22).

**Scheme 1.**
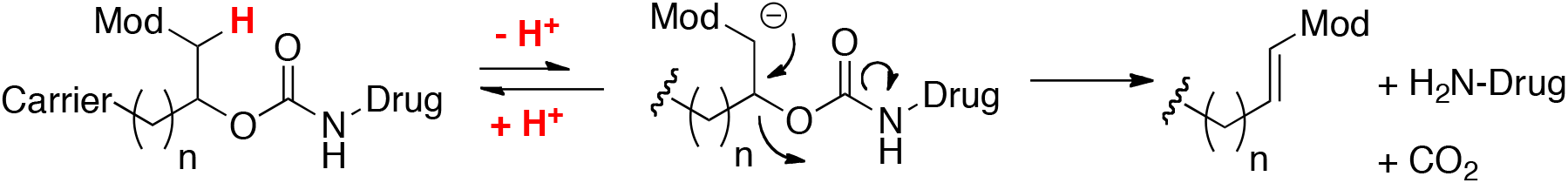
*β*-eliminative drug release from linker

One carrier we use is a mesoporous tetra-PEG hydrogel (23, 24). These hydrogels – fabricated as uniform ∼50 μm microsphere (MS) particles (22, 25) – are deposited locally through a small gauge needle where they slowly release the drug. Slower cleaving β-eliminative linkers are also incorporated in each crosslink of the hydrogel, so gel dissolution occurs in vivo after drug release (26).

Although designed for SC administration, we envisioned that if injected IT, these MS particles would behave as stationary depots that slowly release the drug and bathe the tumor. Indeed, Ascendis’ TransCon technology, with similar objectives but quite different chemistries and attributes, has recently been used for IT administration of a long-acting TLR7/8 agonist (27). Unlike other anchoring approaches, microspheres are not retained by interactions with ECM components, but are rather deposited as stationary particles at the site of injection until they biodegrade. Major advantages of our MS∼drug conjugates for IT injections – as well as peritumoral or intra-nodal injections – are that a) they provide predictable and tunable release rates of the native drug by a simple chemical modification of the linker; b) modification of native proteins is not needed to achieve long IT half-lives, and c) the MS carriers can be used with any class of drugs – peptide, protein or small molecule – with similar behavior.

The primary objective of this work was to establish a generic platform technology to enable long-acting IT therapy. First, using fluorescent dye surrogates, we developed an approach to directly monitor IT pharmacokinetics of MS∼drug conjugates, and showed that IT drug release rates followed the rates expected from the linkers used. Next, we prepared a MS∼small molecule drug conjugate and determined its pharmacokinetic and antitumor effects after IT and systemic administration. Finally, we demonstrate the utility of long-acting IT therapies as a key component of an enabling technology for the safe use of combinations of two drugs with overlapping toxicities.

## Results

### Preparation and characterization of MSs with fluorescent probes

We developed technologies using fluorescent probes that facilitate direct determination of the local pharmacokinetics of drugs released from MS prodrugs (**Fig. 1)**. First, we coupled amine-derivatized microspheres to fluorescein (FL) by a stable linker (FL_s_) to give FL_S_-MS (**1**); here, the FL serves as a quantitative marker for MS particles. We then attached rhodamine piperazine (RP) – a fluorescent drug surrogate – to FL_S_-MS via a releasable β-eliminative linker (RP_R_) to give FL_S_-MS∼RP_R_ (**2**). To measure the release rate of RP, the FL_S_-MS∼RP_R_ and free RP were separated at various times, and the free RP and MS-bound RP_R_ and FLs were quantified. To avoid errors due to inconsistent recovery of MSs, the fluorescence signals of RP_R_ remaining on MSs were normalized to the amount of MS recovered as determined by the FLs content; plots of RP_R_/FL_S_ vs t were used for ratio-metric determinations of release rates of the drug surrogate from the MSs. As described in later experiments where a drug is used instead of the RP_R_ surrogate (e.g. SN-38, vide infra), stable MS-FL_s_ and releasable MS∼drug conjugates were prepared separately and then mixed to give the desired proportions for in vivo experiments.

**Figure 1.**
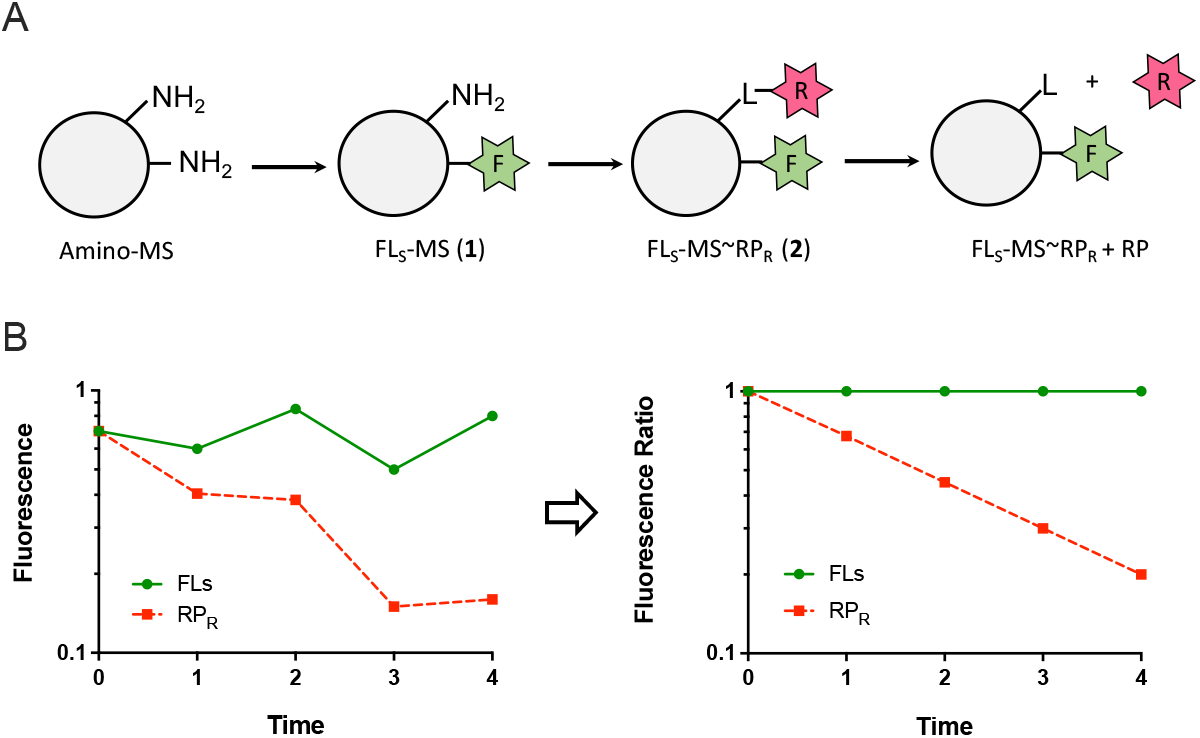
Direct assay for fluorescent IT pharmacokinetic analyses. A) Synthesis of double-labeled microspheres with stable-linked fluorescein (FL_S_) as a quantitative marker for MS particles and releasable rhodamine (RP_R_). B) Depiction of fluorescence measurements with variance in recovery of MS from tissue and the normalized florescence ratio (RP_R_/FL_S_).

The releasable β-eliminative linkers used for RP in FL_S_-MS∼RP_R_ had modulators 4-ClPhSO_2_-(**2A**), PhSO_2_-(**2B**), and 4-MePhSO_2_-(**2C**) (20). The in vitro rates of release of RP from these conjugates (**Fig. S2**) were determined at pH 8.4, 37 °C, and the estimated t_1/2_ values at pH 7.4, 37 °C (28) were 9 hour for **2A**, 27 hour for **2B**, and 40 hour for **2C**. These MS also contained base-labile β-eliminative linkers (Mod = -(CH_3_CH_2_)_2_NSO_2_-α-Lys) within each crosslink to allow solubilization in base prior to analyses; the MSs had a time to reverse gelation (t_RG_) of ∼1000 hours at pH 7.4, 37 °C (22, 25).

### Separation of intact FL_S_-MS from free RP under mock in vivo conditions

We next developed methods to separate intact MS particles labeled with FL (**1**) from free RP under conditions simulating analyses of biopsies of local injections. In early experiments we found that MSs sheared upon sonification, but were stable to mechanical homogenization. When homogenized suspensions containing FL_S_-MS and spiked free RP were centrifuged and the pellet washed, the pellet retained >98% of the FL_S_-MS; the free RP was hardly detectable in the pellet, but present at high levels in the supernatants. In a similar experiment, when the mixture of FL_S_-MS and free RP was passed through a 0.2 μ PTFE spin filter and the retentate washed, the retentate contained >99% of the FL_S_-MS whereas the filtrate contained only released RP (**Fig. S4**). Hence, either centrifugation or filtration followed by washing effectively separates particulate MSs from free fluorescent drug surrogates.

### Half-life of free RP in tissue and normalization of data for ratio-metric analysis

To estimate the t_1/2_ of diffusion of free RP in tissues, a mixture of stable Fl_S_-MS marker and free RP was injected SC in temporal subcutaneous injections at six locations on a rat (26). After euthanasia, tissue samples of 150 ± 22 mg surrounding the injection sites were obtained using a 12 mm biopsy punch, homogenized and analyzed for free RP and Fl_S_-MS. The raw data were normalized to the recovered Fl_S_-MS (**Fig. S5**), and using the ratio-metric method of plotting RP/FLs vs t we calculated a local t_1/2_ of 1.5 ± 0.25 hour for RP in SC tissue.

### Intra-tumoral t_1/2_ values of RP released from FL_S_-MS∼RP_R_

In an exploratory experiment, three groups (n=2) of Balb/c mice were implanted SC with CT26 murine colon carcinoma cells. When tumors reached ∼265 mm^3^, each was injected with 25 μL of FL_S_-MS∼RP_R_ **2A, 2B** or **2C**. After 24 hours, animals were euthanized, tumors and surrounding tissue (150 ± 22 mg) were harvested using a 12 mm biopsy punch. Samples were mechanically homogenized, pellets were obtained by the centrifugation method and thoroughly washed. After hydroxide dissolution of the MSs, fluorescence of FL_S_ and RP_R_ were quantified. **Fig. 2A** shows the data at 24 hours for each dye in conjugates **2A, 2B**, and **2C** normalized to the amount of FL_S_ in the pellet. There is a high level of FL_S_ in the pellets – a measure of the isolated microsphere depot – with <1.6% in the extracts. Most RP was also found in the pelleted FL_S_-MS∼RP_R_, and the amounts track the t_1/2_ of the linkers in the order **2A**<**2B**<**2C**, indicating that pelleted RP emanates from FL_S_-MS∼RP_R_ and not residual free RP.

**Figure 2.**
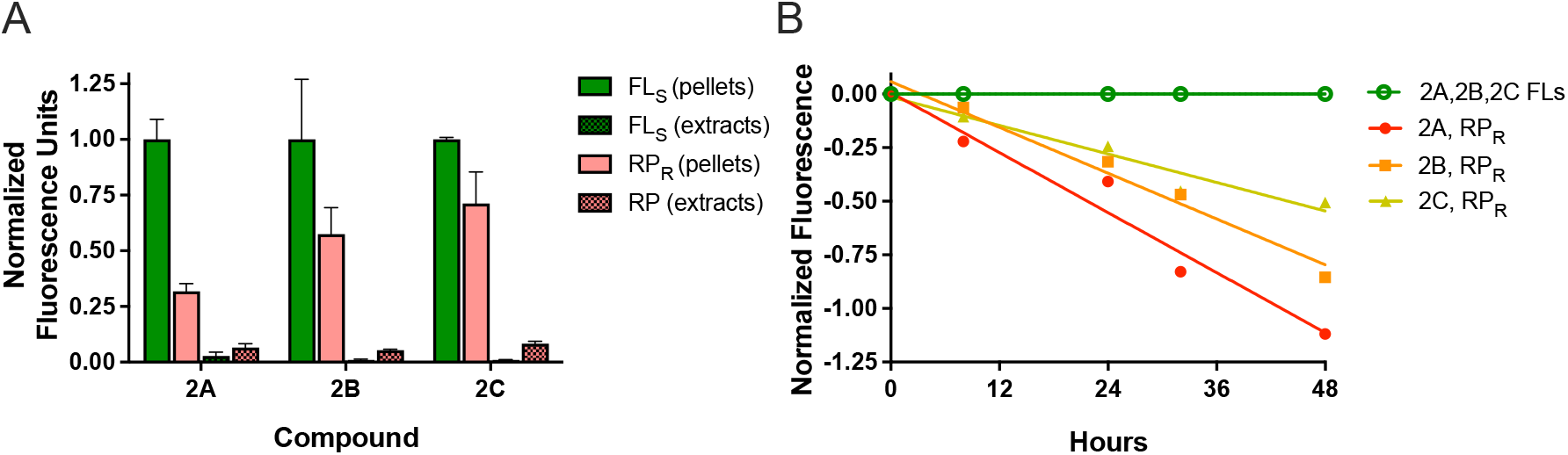
Separation of MSs and released drug surrogate 24 hour after IT injection and in vivo IT pharmacokinetics. A) Normalized fluorescence of FL_S_-MS∼RP_R_ and free RP 24 hour after injection into tumors. B) Normalized fluorescence of C vs t plots of **2A, 2B** and **2C** after IT injections of FL_S_-MS∼RP_R_. Normalization was performed as follows: RP = Ln[(RP/FLs)/(RP/FLs)t0)] and FLs = Ln(FLs/FLs).

We performed a larger experiment in which murine CT26 colon carcinoma tumors (n=15) were injected IT with MSs **2A, 2B** and **2C**. Three tumors containing each conjugate were harvested at 0, 8, 24, 32 and 48 hours and the isolated MSs were analyzed as above for FL_S_ and RP_R_ content over 48 hr. **Fig. 2B** shows the normalized ratio-metric plots of these experiments from which we determined t_1/2_ values of 30 hour for **2A**, 43 hour for **2B** and 63 hour for **2C**. Together, these results show that the IT t_1/2_ of the surrogate drug released from the MSs is governed by the linker used.

### Release rates in subcutaneous/peritumoral tissues

Because drug release from the MSs is rate-determining the tumor t_1/2_ of the drug should be similar in different tissues – SC, peritumoral and IT – unless there is a direct effect on the linker cleavage. When **2C** (Mod PhSO_2_-) was injected SC rather than IT the release t_1/2_ was 43 h, some 1.5-fold faster than the 63 hour t_1/2_ when injected IT. Drug release from MS conjugates occurs by a first-order base catalyzed elimination, and the tumor pH is lower than the peritumoral/SC pH (29). Hence, the slower release rate of IT vs SC injected MS∼RP is simply explained by the lower pH in tumors which slows linker cleavage.

### Preparation and characterization of MS∼SN-38

Microspheres with releasable SN-38 (MS∼SN-38) were synthesized using methods adapted from the above method and Schneider et al. (28). Briefly, azido-linker(SO_2_Me)-SN-38 (30) was attached to bicyclooctyne (BCN) derivatized microspheres by SPAAC, generating MS∼SN-38 (**Scheme 2**). The MS∼SN-38 were extensively washed, then centrifuged to give a slurry of MS∼SN-38. HPLC analyses of the supernatant indicated there was no residual N_3_-linker(SO_2_Me)-SN-38 or SN-38 in the packed slurry. The extent of loading of SN-38 on the MS was determined by dissolving 20 μL of the slurry in 50 mM NaOH and quantifying SN-38 by A_414_, and PEG by a colorimetric assay (22). The SN-38 loading was 196 nmol SN-38/mg PEG or ∼5 μmol/mL slurry, 98% of the maximum. The in vitro t_1/2_ of release of SN-38 from MS∼SN-38 conjugates was ∼160 hour at pH 7.4, 37°C (**Fig. S7**) and the SN-38 released at pH 8.4 was >95% pure.

**Scheme 2.**
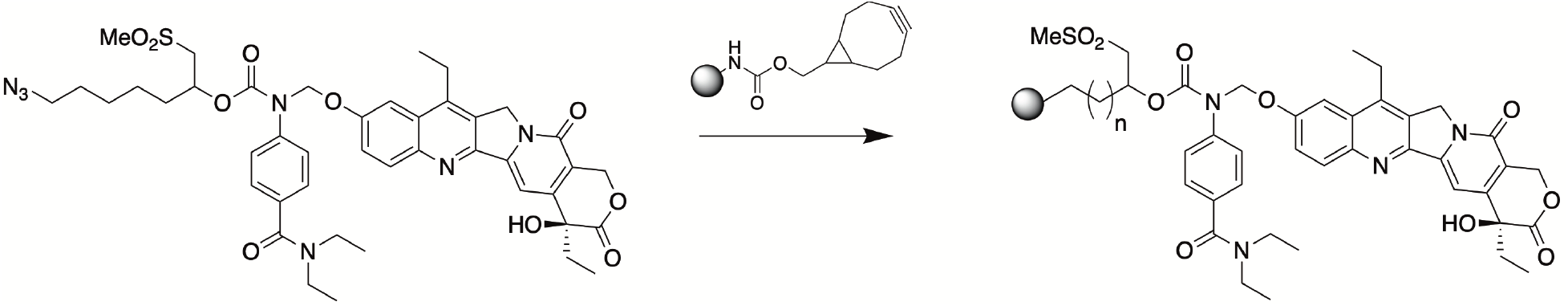
SPAAC reaction to prepare MS∼SN-38

### Release of SN-38 from MS∼SN-38 injected SC in rats

Stable MS-FL_s_ and releasable MS∼SN-38 conjugates were made separately and mixed to give FL_s_-MS∼SN-38 (vide supra); as above, FL_s_ serves as a quantitative marker of the MSs to allow ratio-metric analysis from normalized measurements of SN-38/FL_S_. A 100 uL mixture containing MS∼SN-38 (10 nmol) and MS-FL_s_ (32 μM FL) was administered SC to four locations on the backs of seven SD rats. At specified times each rat was euthanized and samples containing the MSs and surrounding tissue were excised with a 12 mm diameter biopsy punch. Each of the tissue samples was dounced in 0.5% HOAc, clarified by centrifugation, and washed with 0.5% HOAc. Then the pellets were treated with 0.25 M NaOH to dissolve MS∼SN-38. The AcOH extracts and digested pellets were each analyzed by HPLC for free SN-38 and FL_s_.

A plot of the free SN-38 in the AcOH extracts and NaOH tissue pellets normalized to the fluorescein content of the MSs (i.e. SN-38/FL_S_) vs t gave a t_1/2_ of 6.6 days (R^2^ of 0.974) for the free SN-38 in the extract, and 6.1 days (R^2^ 0.910) for the SN-38 in the pellet (**Fig. 3**). Correcting for the 1.5 slower t_1/2_ of linker cleavage in tumor vs SC tissue (vida supra), the IT t_1/2_ in CT26 tumors is estimated to be ∼10 d.

**Figure 3.**
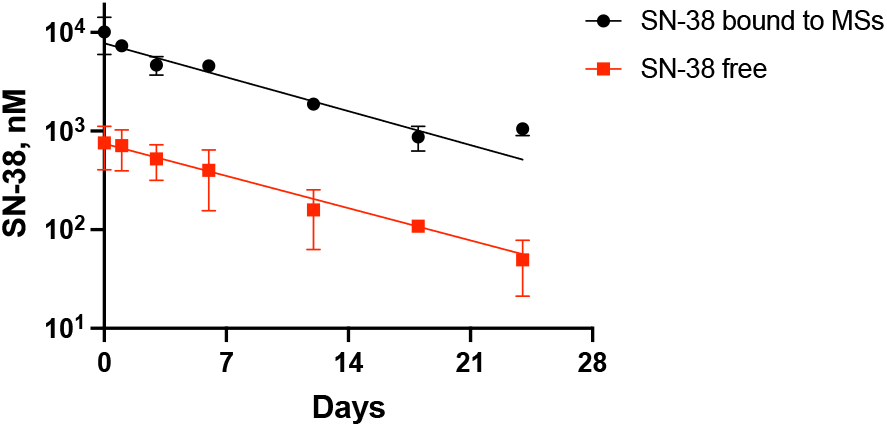
MS∼SN-38 and free SN-38 recovered from MS containing skin plugs over time. Data normalized to the fluorescein marker for MS particles. Error bars represent one standard deviation of the mean (n=4/point).

### Tumor growth inhibition by IT vs SC MS∼SN-38

We wanted to compare the tumor growth inhibition (TGI) of IT SN-38 released from IT MS∼SN-38 to systemic SN-38 derived from PEG∼SN-38 and SC MS∼SN-38. We initially compared the effects of 7.5 μmol/kg IT MS∼SN-38 to equimolar IP PLX038A, a highly effective PEG∼SN-38 prodrug on 22Rv1 ATM^-/-^ tumors, chosen because it is sensitive to both TOP1i and PARPi (**Fig. S8**) (31). The MS∼SN-38 showed two-fold longer tumor suppression and survival time of compared to the PEG∼SN-38 prodrug.

**Fig 4A,B** shows tumor growth and host survival in mice (n=5) bearing subcutaneous 22Rv1 ATM^-/-^ tumors after a single IT injection of MS∼SN-38 at 0.2- to 7.5 μmol/kg (38-150 nmol/mouse). The lowest IT dose of 0.2 μmol/kg resulted in a TGI ∼55% and 13 day extension in overall survival; the highest 7.5 μmol/kg dose resulted in ∼90% TGI and 41 day extension in overall survival. **Fig 4C,D** also shows tumor growth and host survival after SC injection of MS∼SN-38 at 0.2- and 2.0 μmol/kg. Whereas the low dose gave insignificant TGI, the 2.0 μmol/kg gave 50% TGI and survival, comparable to 0.2 μmol/kg IT MS∼SN-38. We also determined that IT MS∼SN-38 could control growth of very large tumors. When 22Rv1 ATM^-/-^ tumors reached an average tumor volume ∼1,000 mm^3^, 2- or 20 μmol/kg MS∼SN-38 was administered IT. Although 2 μmol/kg MS∼SN-38 resulted in a modest ∼1 week delay in tumor growth and a 15 day increase in median survival, IT administration of 20 μmol/kg MS∼SN-38 resulted in >42 day delay in tumor growth and a median survival of 45 days (**Fig. 4E,F)**. Hence, IT MS∼SN-38 is about 10-fold more potent than systemic administration and can suppress growth of very large tumors.

**Figure 4.**
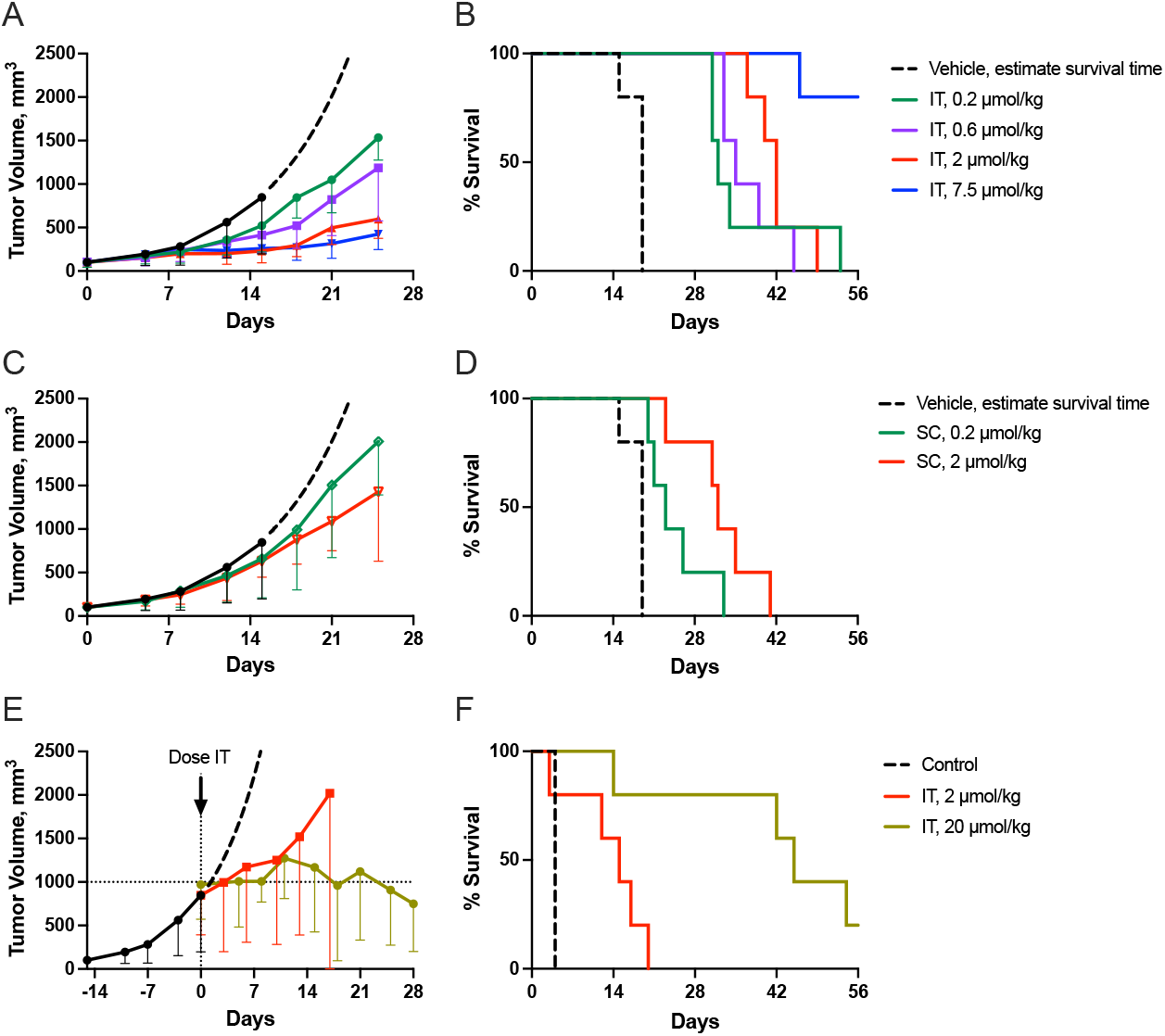
Anti-tumor effects of MS∼SN-38 in mice bearing 22Rv1 ATM^(-/-)^ tumors. **A)** Tumor volume vs days post treatment; control (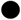) showing estimated exponential growth after 1,000 mm^3^ (**– –**); IT MS∼SN-38 doses were 0.2-(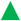), 0.6-(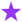), 2-(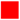) or 7.5 μmol/kg (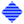). **B)** Kaplan Meier plot of survival for animals treated IT with 0.2- to 7.5 μmol/kg MS∼SN-38. **C, D)** Tumor volume vs days post treatment and corresponding Kaplan Meier plot of overall survival following SC 0.2-(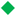) and 2 (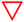) μmol/kg MS∼SN-38. **E, F)** Tumor volume vs time post treatment and corresponding Kaplan Meier plot of survival of mice with 1,000 mm^3^ tumors treated with 2 - (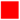) or 20 μmol/kg (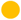) IT MS∼SN-38. Data show median tumor volume ± median absolute difference from the median (n=5/group). Survival times of the controls (**– –**) are based on the estimated times for the tumor volume to reach 2,000 mm^3^. The estimated median survival time of the control group was 19 days.

### Tumor inhibition by IT MS∼SN-38 in combination with systemic talazoparib (TLZ)

TOP1i and PARPi are highly synergistic towards BRCA and ATM deficient tumors, but have overlapping hematologic toxicities (32, 33). We wanted to know if SN-38 that was localized in a tumor by IT MS∼SN-38 in combination with systemic administration of a PARPi would have synergistic anti-tumor effects. We measured the effects of a combination of IT MS∼SN-38 with the PARPi TLZ on growth of TOP!i and PARPi sensitive 22Rv1 ATM^-/-^ xenografts. Tumor growth (TG) and tumor growth inhibition (TGI) were calculated as described (34) using the fractional AUC for the treated group in relation to that of the vehicle. We assessed the interaction of the drug combination using an additivity index determined as TG_calc_/TG_obsd_ where TG_calc_ is the product of TG values of the individual drugs and TG_obsd_ is the observed tumor growth in the presence of both drugs. Here, an index <1 indicates infra-additive, an index of 1 indicates additivity and an index >1 indicates a supra-additive or synergistic interaction. **Fig. 5** shows the growth of 22Rv1 ATM^-/-^ xenografts in mice (n=6) treated with 0.2 μmol/kg MS∼SN-38 as a single IT dose, 0.4 μmol (0.15 mg)/kg of TLZ PO QD, and a combination of both. The TG was 0.33 for TLZ and 0.38 for IT MS∼SN-38 giving a TG_calc_ for the combination of 0.12. The TG_obsd_ of the combination was 0.04 so the additivity index was estimated as 3.0 (0.12/0.04), indicating very strong synergy of the two drugs.

**Figure 5.**
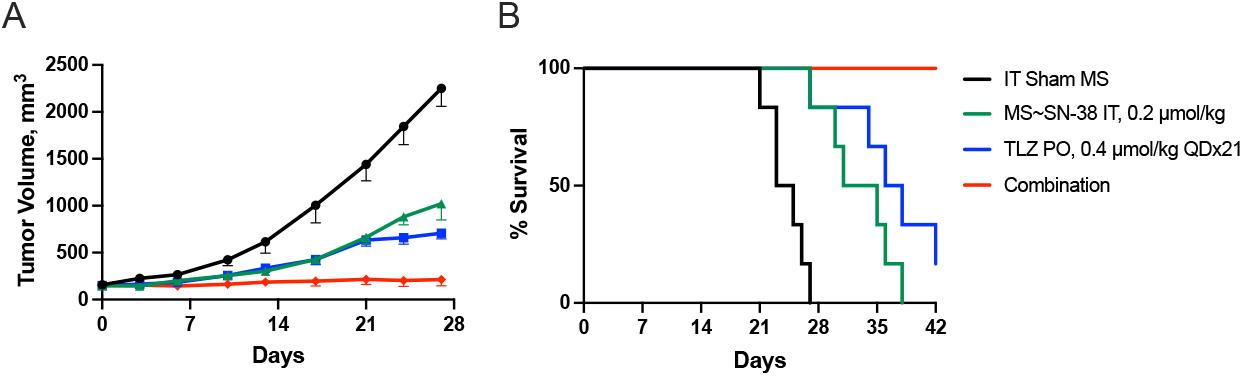
Anti-tumor effect of single dose IT MS∼SN-38 in combination with QD oral TLZ on 22Rv1 ATM^(-/-)^ tumors. **A)** Median tumor volume vs time post treatment; animals received IT injection of empty MSs (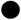), 0.2 μmol/kg IT MS∼SN-38 (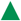), 0.4 μmol/kg TLZ PO QDx21 (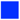), or the combination of 0.4 μmol/kg TLZ PO QDx21 and 0.2 μmol/kg IT MS∼SN-38 (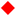). Data are median tumor volume ± median absolute difference from the median (n=6/group). B) Kaplan Meier plot of overall survival. The median survival times for the vehicle control (24 d), 0.2 μmol/kg IT MS∼SN-38 (33 d), 0.4 μmol/kg TLZ PO QDx21 (37 d), and the combination (>42 d) are based on the tumor volume reaching 2,000 mm^3^ or animal death.

## Discussion

IT therapy has assumed an important role in delivery of immuno- and chemotherapeutics (1, 2, 35). Between January 2018 and June 2021 there were 153 IT clinical trials initiated (36) and a single institution performed over 1,500 image-guided IT immunotherapy injections over a few years (6). The significant advantage of IT injection is that low doses can achieve extraordinarily high IT drug concentrations in the setting of very low systemic exposure, resulting in high efficacy and low toxicity. Notwithstanding proven benefits, IT therapy has not yet realized its promise or acceptance as a foremost approach for cancer therapy.

Although drugs administered IT ultimately enter the systemic circulation, they initially transit through the tumor at rates governed by tumor transport and drug properties (37). An underappreciated problem with IT injections is the rapidity with which drugs leak out of a tumor, leading to reduced exposure and a need for impractical frequent injections (10, 38). For example, IT residence of a protein therapeutic is a function of size (9) but even large mAbs have an IT t_1/2_ of only ∼6 hour (7, 8), hardly sufficient to achieve high exposures for maximal efficacy. Indeed,

we and many others believe that the major reason IT – as well as peritumoral or intra-nodal injections – have not been more successful is their short local residence time.

Half-life extension of IT therapeutics greatly increases their anti-tumor efficacy and favorable effects on immune responses (7, 9, 10, 39). For example, Wang et al. (39) showed much superior efficacy of IT IL-15, PD-1 and combinations of the two when they administered them as a long-acting hydrogel with a ∼7 day t_1/2_. Likewise, Momin et al. (10) showed that extending the IT t_1/2_ of locally-injected IL-2 by increasing molecular size and improving matrix-affinity increases therapeutic efficacy in mice. Of the existing methods for achieving IT t_1/2_ extension, the most efficacious involve some form of “anchoring” to the site of injection, usually through binding of the therapeutic to ECM components (9, 10, 15).

We previously developed and have extensively described (See refs. (21, 40), https://www.prolynxinc.com) a unique technology for long-acting SC injections. Here, a drug is covalently tethered to long-lived hydrogel MSs by a cleavable linker. After SC injection, the particulate MSs deposit locally and the linker slowly cleaves to release the native drug which then enters the systemic circulation. Providing that drug release from the MSs is slower than its systemic elimination, when a MS∼drug is delivered SC the local half-life of the released drug mirrors the half-life determined from analysis of plasma pharmacokinetics; thus, substantial information on the interstitial half-life of such drugs already exists. Also, the t_1/2_ of the drug should be similar in different tissues – SC, peritumoral and IT – unless there is a direct effect on the rate-determining linker cleavage. Since the only determinants of in vivo linker cleavage rates are the modulator used, the pH and the temperature (20), peritumoral and IT cleavage rates can only differ from SC rates because of lower pH. Although the lowered tumor pH will reduce the rate of cleavage, the slower rate may be beneficial because it provides longer drug exposure. In the present work, we show that the same system used for SC administration provides a superior generic delivery system for IT therapeutics. That is, the particulate MSs deposited IT slowly release the attached drug to “bathe” the tumor for long periods before it diffuses to the systemic circulation.

This approach has several advantages over other technologies for IT half-life extension. A major advantage is the unique ability to tune the rate of drug release by simply modifying the linker. Thus, tunable IT pharmacokinetics can be exploited to optimize exposure of the targeted tumor. It also follows that, unlike other anchoring technologies, MS carriers could be used with any class of drugs – peptide, protein or small molecules – and give similar pharmacokinetic behavior; indeed, we have demonstrated this with numerous MS conjugates; see for e.g. (21, 24, 40).

Another benefit is the very long local half-lives that can be practically obtained. As examples, with this technology the half-life of a short-acting 39 AA peptide was extended from ∼30 min to 1 month (21) and that of a cyclic octapeptide was extended to over 2 months (41). The only other reported technology that can achieve this duration of exposure is the alum-tethering approach, but this is limited to proteins and requires their structural modification (18, 19).

Also, because the MS particles are physically deposited in or around the tumor before drug release, they have advantages over approaches for IT delivery that use drug solutions. First, unlike most anchoring approaches they do not require a tumor-localization domain or an ECM-binding target protein. Second, MS particles would avoid the problem of drug “spillover” that occurs with volumes of soluble drugs over the 25- to 50% “hold up” volume of solid tumors (10, 42, 43). Last, they are not affected by biophysical barriers within the tumor (44) that can result in drug leakage with consequent dose variability and systemic toxicity. Hence, the MS∼drug conjugates should provide a simple solution to some major problems in IT therapy.

The present work was undertaken to establish a generic platform technology that enables long-acting IT therapy. The specific objectives were to 1) develop technology to directly monitor IT pharmacokinetics of MS∼drug conjugates, 2) determine the pharmacokinetics and antitumor effects of IT MS∼SN-38 conjugates, and 3) demonstrate utility of the technology to enable safe use of combinations of two drugs with overlapping toxicities.

First, we developed approaches for direct determination of IT pharmacokinetics in tissue that, in principle, could be adapted for human biopsy specimens. After showing that MSs and free drug in tissue biopsies could be cleanly separated, we developed methods that allow quantitation of the free drug, as well as the drug remaining on recovered MSs. Here, MSs were labeled with fluorescein by a stable linker to serve as a quantitative marker for particles; MSs were also prepared in which rhodamine was attached by a releasable linker to serve as a surrogate for a releasable drug. When combined, the rhodamine/fluorescein ratio on the MSs measure the amount of rhodamine remaining on the MSs, and normalizes the measurement independent of the efficiency of particle recovery. With this approach, we confirmed that the IT release rate of RP from MS∼RP_R_ conjugates with different linkers correlated to their in vitro release rates. Using the same fluorescein quantitative marker for particles, the ratio-metric method can be adapted to determination of the IT pharmacokinetics of MSs attached to non-fluorescent drugs.

Next, we determined the local pharmacokinetics and anti-tumor effects of IT MS∼SN-38 conjugates. SN-38 is a potent inhibitor of topoisomerase 1 (TOP1) that causes single- and double stranded DNA breaks which initiate the DNA damage response (DDR) (45, 46). TOP1i also have immune modulatory effects including activation of the STING pathway (47), sensitization of tumors to checkpoint inhibitors (48) and immunogenic cell death (49, 50). In BRCA- and ATM-deficient tumors the DDR is lacking important repair enzymes that makes the tumors particularly sensitive to DNA damage – as by a TOP1i – as well as inhibitors of DDR enzymes – such as PARPi (46, 51). We therefore studied the effects of MS∼SN-38 on BRCA and ATM deficient, TOP1i-sensitive 22Rv1 xenografts.

We attached SN-38 to MSs using a linker with a MeSO_2_ modulator to give a MS∼SN-38 conjugate with an in vitro release t_1/2,pH 7.4_ of 150 h. When MS∼SN-38 containing 10 nmol SN-38 was administered SC, the normalized tissue C vs t plots showed that in vivo release of SN-38 from the MSs occurred with a t_1/2_ of ∼6.5 d, very close to the in vitro t_1/2_ of ∼6 d. Importantly, the released free SN-38 reached a very high C_max_ of ∼1 μM, and remained above 10 nM – a concentration sufficient for TOP1 inhibition – for ∼3 Wk. After correcting for the slower t_1/2_ of linker cleavage in the acidic tumor vs neutral SC tissue, the IT t_1/2_ in CT26 tumors was estimated as ∼10 d.

To determine antitumor effects of IT vs systemic delivery of SN-38, tumor growth inhibition and host survival of TOP1i-sensitive (32) 22Rv1^ATM-^ xenografts were determined after treatment with various concentrations of IT and SC MS∼SN-38. Both routes of administration ultimately deliver SN-38 systemically to the same extent, but after IT injection high concentrations of the released SN-38 transit through the tumor tissue before entering the systemic circulation. We reasoned that if the efficacy of IT SN-38 was due to systemic exposure the IT and SC dosing required for efficacy should be the same; however, if the efficacy of IT administration was due to a local effect, the IT injection would be more potent. Gratifyingly, IT administration of MS∼SN-38 required a ten-fold lower dose than SC administration to achieve the same tumor growth inhibition and host survival. Taken together, these results show that the anti-tumor effect IT MS∼SN-38 must be due to a local vs systemic effect, and IT administration of MS∼SN-38 is ∼10-fold more effective than systemic administration.

Finally, we demonstrated that the ability to localize a drug in a tumor for prolonged periods may enable an approach for the safe use of a combination of drugs that have overlapping toxicities – which is very common in cancer chemotherapy. In this approach the long-acting IT drug is injected into the tumor along with systemic treatment with the second drug. Here, the tumor is exposed to the combined effects of both drugs, but all other tissues may be exposed only to the single systemic drug. Hence, the drugs may show additive or synergistic effects in tumor inhibition, but tissues may only have toxic effects due to the single systemically administered drug.

In the present context, TOP1i and PARPi are highly synergistic in growth inhibition of tumors but have overlapping myelosuppression that prevents safe use of the combination (33, 46). Here, we wanted to know if SN-38 localized in a tumor by IT MS∼SN-38 would have synergistic anti-tumor effects in combination with systemic administration of a PARPi. When a single low dose of MS∼SN-38 was administered IT to 22Rv1^ATM-^ xenografts and oral TLZ was administered QD, high anti-tumor synergy of the combination was observed. Since the same amount of single-agent SN-38 administered systemically has little effect, we surmise that there is a low probability of overlapping systemic toxicity. We posit that this approach to safely combining two drugs with overlapping toxicities might be executed with MS∼drug conjugates of other IT therapeutics.

Indeed, there are many potential uses of this technology for safely administering combinations of immuno-oncology agents. For example, immune checkpoint inhibitors (ICI) such as ***α***CTLA4 and ***α***PD-1 and their combination have shown remarkable activity against multiple tumor types, and can lead to durable remissions (52). However, the success of ICI has been limited by inflammatory immune-related adverse events (irAEs). Combination of both ***α***CTLA4 and ***α***PD-1 – which has been approved for several cancers – is much more effective than either single agent but it is also considerably more toxic (53). A possible approach to treat a tumor with both agents and avoid irAE-associated toxicities would be to treat the tumor with a long-acting IT ***α***CTLA4 along with systemic ***α***PD-1. The tumor could be exposed to the synergistic effects of both drugs but the irAEs in normal organs would be limited to those of single agent systemic ***α***PD-1.

A limitation of this work is that it was solely directed towards xenografts of human tumors in immunodeficient mice and do not address effects the immune system might have on the efficacy of the drug treatments. The intent was to develop a technology for long-acting IT therapy, and an approach to safely deliver therapies with overlapping toxicities in human cancer cells in vivo. Likely, because TOP1i have significant immune modulatory effects (46-49), future studies in immunocompetent mice may uncover aspects of long-acting IT delivery of the agents that better reflect expectations in patients.

In conclusion, the releasable MS∼drug technology – originally developed for long-acting SC administration – serves well as a generic technology for long-acting IT therapy. In addition to the results on small molecules described here, we have also shown the technology is effective in increasing the IT half-life of IgGs in NSG mice from 1.5 day to ∼ 1 month (unpublished results) which makes it amenable for IT use with mAb immuno-oncology agents, antibody drug conjugates and other large proteins. We describe an approach for direct analysis of tissue and tumor pharmacokinetics of MS∼drug prodrugs, and the favorable tissue pharmacokinetics of the DNA damaging agent SN-38 delivered by a MS∼SN-38 conjugate. We have also shown that IT injection of MS∼SN-38 in 22Rv1^ATM-^ xenografts has anti-tumor activity comparable to a 10-fold higher dose of systemically administered SN-38. Finally, we describe and demonstrate how long-acting IT injections might enable safe use of drug combinations with overlapping toxicities. The long-acting IT drug – here, MS∼SN-38 – is administered to the tumor and the second drug – here, a PARPi – is administered systemically. Since the tumor is exposed to both drugs while all other tissues are only exposed to the systemic drug, selective toxicity of the tumor might be achieved. We posit that variations of this approach provides a generic approach for safe use of drug and immunotherapy combinations with overlapping toxicities.

## Materials and Methods

The sources of specialized materials are provided along with their use in *SI Appendix*. Detailed synthetic, conjugation, and analytical procedures are described. In vitro kinetic procedures are provided as are in vivo pharmacokinetic and pharmacodynamic methods and analyses. All animal handling and care was performed by MuriGenics (Vallejo, California) and conformed to IACUC recommendations.

## Data Availability

All study data are included in the article and/or *SI Appendix*.

## Acknowledgements

DVS wishes to thank T.J. Vindigni for helping with construction of the manuscript, and R. Reid for assistance on modeling pharmacokinetics.

## Author contributions

J.H., J.A.H., D.C. conceived project, performed experiments, interpreted results; C.W.C. performed experiments; G.W.A., D.V.S. conceived project, interpreted results; D.V.S. wrote manuscript.

