## Supplemental Information for "Releasable hydrogel microsphere-drug conjugates as generic prodrugs for long-acting intra-tumoral therapy"

**Supplementary Information**

**Table of contents**

1. General Methods

2. Fluorescein labeled microspheres ($FL_s$-MS-Ac)

3. Dual labeled MSs with fluorescein and releasable RP ($FL_s$-MS~$RP_R$)

    3.1 Rhodamine piperazine (RP)
    3.2 Cleavable linkers (azido-linker(Mod)-HSCs)
    3.3 Synthesis of azido-linker-RP conjugates
    3.4 Fluorescein labeled and cyclooctyne activated MSs ($FL_S$-MS-CO)
    3.5 Dual labeled microspheres ($FL_s$-MS~$RP_R$ **2A**, **2B**, and **2C**)
    3.6 *In vitro* RP release rates from $FL_s$-MS~$RP_R$
    3.7 Dosing syringes, dosing accuracy, and $FL_s$/RP ratio for $FL_s$-MS~$RP_R$

4. Assay development, and intra-tumoral injections of $FL_s$-MS~$RP_R$

    4.1 *In vitro* separation of RP from $FL_s$-MS following blending
    4.2 Clearance half-life of RP in tissue, following SC injection into rats
    4.3 Determination of $FL_S$ and RP in tissue
    4.4 Tumor xenografts in mice and intra-tumoral injection of $FL_s$-MS~$RP_R$
    4.5 Determination of $FL_S$ and RP in excised tumors

5. Release of SN-38 from subcutaneous microspheres

    5.1 Preparation of microspheres with releasable SN-38 (MS~SN-38)
    5.2 *In vitro* release and dissolution of MS~SN-38
    5.3 *In vivo* release of SN-38 from microspheres
        5.3.1 Dose formulation preparation and analysis
        5.3.2 Dose administration and recovery
        5.3.3 Analysis of free SN-38 and MS-bound SN-38 at injection sites
    5.4 Anti-tumor efficacy of MS~SN-38
    5.5 Animal Welfare Statement

6. References

### 1. **General Methods**

All reagents were reagent grade. HPLC analyses were performed on a Shimadzu LC-20AD HPLC system equipped with an SPD-M20A diode array detector, a RF-10AXL fluorescence detector, and a Phenomenex Jupiter 5 µm C18 column (300 Å, 150 x 4.6 mm). Unless otherwise noted, peaks were eluted with a 10 minute linear gradient of 20-100% acetonitrile in water (0.1% TFA) at 1 mL/min. Fluorescence measurements were made using a Molecular Devices, Spectramax i3 plate reader, and the following excitation and emission wavelenghts: $FL_S$ ex 485 nm em 535 nm, RP ex 565 nm, em 589 nm. The following extinction coefficients were used for quantification by absorbance: FL $A_{495}$ ($\varepsilon$ = 80,000 $M^{-1}cm^{-1}$ for pH > 9), RP $A_{565}$ ($\varepsilon$ = 96,100 $M^{-1}cm^{-1}$), and SN-38 $A_{363}$ ($\varepsilon$ = 27,500 $M^{-1}cm^{-1}$) or $A_{414}$ ($\varepsilon$ = 27,500 $M^{-1}cm^{-1}$).

### 2. **Fluorescein labeled microspheres ($FL_s$-MS-Ac)**

Stable linked fluorescein labeled tetra-PEG hydrogel MSs with acylated amines ($FL_s$-MS-Ac) containing the –CN α-Lys crosslink cleavage rate modulator were produced from amino-MS and suspended in isotonic acetate tween buffer (IAT, 10 mM pH 5 acetate, 145 mM NaCl, % tween 20) as previously described (1). The fluorescein concentration of the suspension (0.8 mM) was determined by absorbance by dissolving 0.050 mL of suspension into 0.450 mL of 50 mM NaOH then measuring absorbance at $A_{495.}$ These MS were used in experiments when FLs-MS were mixed with other MS-drug conjugates to serve as a tracer for MS at the injection site.

### 3. **Dual labeled MSs with stable fluorescein and releasable RP ($FL_s$-MS~$RP_R$)**

#### 3.1 Rhodamine piperazine (RP)

**Scheme S1.** Synthesis of RP from rhodamine B.

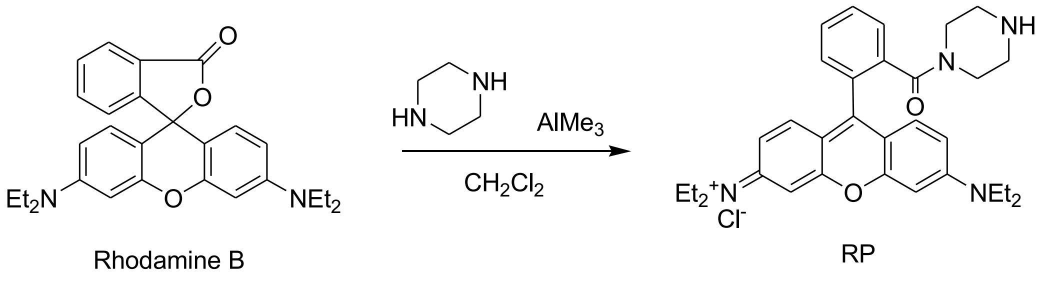


RP was synthesized from Rhodamine B as previously reported (2).

#### 3.2 Cleavable linkers (azido-linker(Mod)-HSCs)

The following azido linker hydroxy succinimidyl carbonates (azido-linker(Mod)-HSCs) with cleavage rate modulating groups: A = $ClPhSO_2$-, B = $PhSO_2$-, and C = $MePhSO_2$- were prepared as previously described (3).

A: O-[1-(phenylsulfonyl)-7-azido-2-heptyl]-O′-succinimidyl carbonate

B: O-[1-(4-chlorophenylsulfonyl)-7-azido-2-heptyl]-O′-succinimidyl carbonate
C: O-[1-(4-methoxyphenylsulfonyl)-7-azido-2-heptyl]-O′-succinimidyl carbonate

3.3 Synthesis of azido-linker-RP conjugates

**Scheme S2.** Synthesis of azido-linker-RP conjugate A.

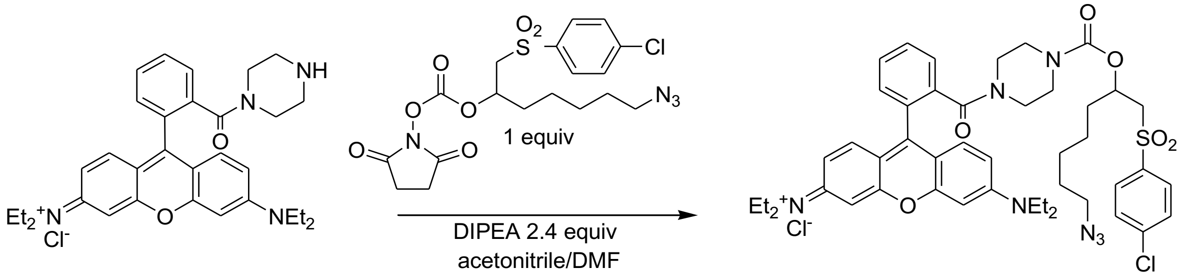


The following procedure (**Scheme S2**) was used to synthesize azido-linker-RP conjugates A, B, and C from azido-linker(Mod)-HSCs A, B, and C. Azido-linker(Mod)-HSC (0.0075 mmol, 1 equiv) was treated with solution of RP (3.8 mg, 0.0075 mmol, 1 equiv) and DIPEA (2.3 mg, 0.018 mmol, 2.4 equiv) in 1 mL of acetonitrile containing 20% (v/v) DMF. After 2 h the reaction mixtures were analyzed by reverse phase HPLC with detection at 261 nm. All reactions showed greater than a 95% yield of the azido-linker-RP conjugate, as well as the presence of some unreacted rhodamine B that was present in the starting material (**Fig. S1**). The reaction mixtures were then used as is for loading of the azido-linker-RP conjugates onto MSs as described below.

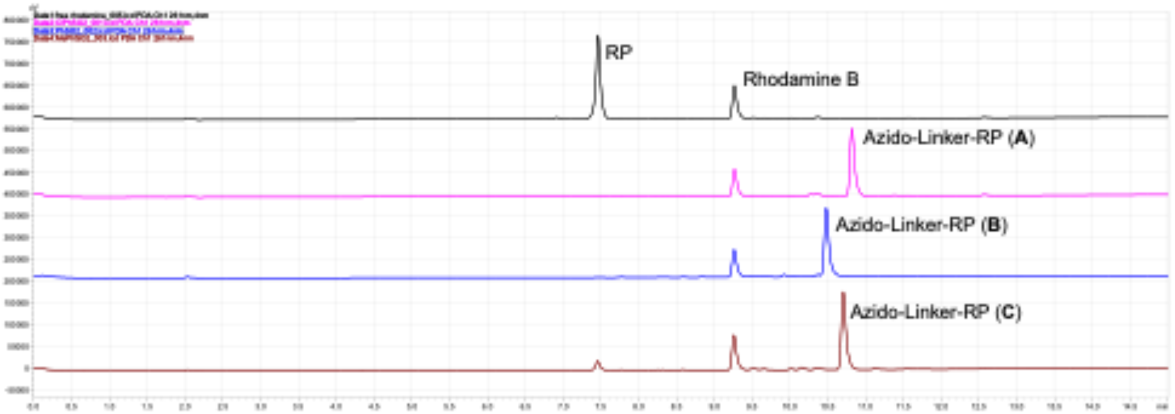


**Figure S1.** C18-HPLC analysis of reaction mixtures, showing RP, and Azido-linker-RP A, B, and C.

3.4 Fluorescein labeled and cyclooctyne activated MSs ($FL_S$-MS-CO)

**Scheme S3.** Synthesis of $FL_S$-MS-CO and $FL_S$-MS~$RP_R$ from amino-MS.

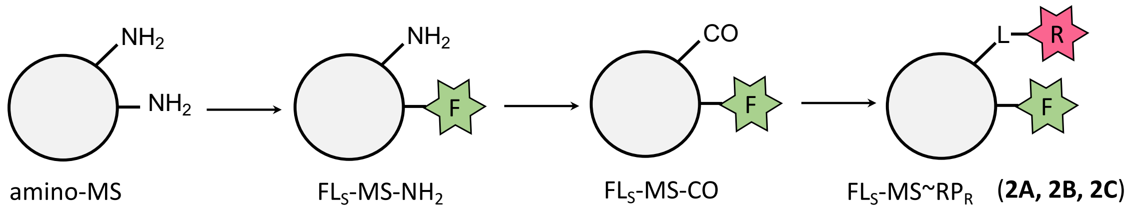


A slurry of amino-MS containing the $(CH_3CH_2)_2NSO_2$- α-Lys crosslink cleavage rate modulator (4) in acetonitrile (45 mL, 3.3 mM $-NH_2$, 0.150 mmol $-NH_2$) was treated with DIPEA (76.8 mg, 0.104 mL, 0.594 mmol, 4 equiv) in acetonitrile (10 mL) and a solution of (5-(and-6)-carboxyfluorescein succinimidyl ester (Invitrogen C1311, 14 mg, 0.0297 mmol, 0.2 equiv) in 0.8 mL DMF. After 35 min at room temperature a solution of 5-HCO-OSu (5) (1 mL, 150 mM, 0.150 mmol, 1 equiv) in acetonitrile was added. After standing 1.5 h, the slurry was washed three times with acetonitrile (80 mL), using centrifugation at 4500 x G between washes to separate the MSs from supernatant. The slurry was then divided into five equal portions in 50 mL centrifuge tubes (Falcon). Each tube contained ~8 mL of packed $FL_S$-MS-CO (**Scheme S3**), containing 0.024 mmol of cyclooctyne, and was used as is for attachment to azido-linker-RP conjugates.

3.5 Dual labeled microspheres ($FL_s$-MS~$RP_R$ **2A**, **2B** and **2C**)

The following procedure (**Scheme S3**) was used to synthesize azido-linker-RP MS conjugates **2A**, **2B**, and **2C**. A slurry of $FL_S$-MS-CO (0.024 mmol CO, 1 equiv) in acetonitrile (8 mL) was treated with one of the azido-linker-RP containing reaction mixtures (A, B, or C, 1 mL, 0.0075 mmol, azido-linker-RP, 0.3 equiv), methanol (5 mL), and acetic acid (0.001 mL). The resulting slurries were kept at room temperature for 36 h then centrifuged at 4500 x G to separate the MSs from supernatant. The MSs were then washed with 1 x 40 mL of methanol, separated by centrifugation at 4500 x G then treated with a solution of O-(2-Azidoethyl)heptaethylene glycol (0.5 mL, 40 mM, 0.020 mmol, 0.8 equiv) in water, to cap unreacted cyclooctynes. After 18 h the MSs were washed with 3 x 40 mL methanol, then with 4 x 40 mL of IAT buffer centrifuging at 4500 x G in between washes. Finally the $FL_s$-MS~$RP_R$ slurries were transferred to 10 mL syringes and packed at 3000 x G for 5 min using a special fixture for centrifugation of the syringes (6) to provide ~5 mL of $FL_s$-MS~$RP_R$ slurry for each material (**2A**, **2B**, and **2C**). Measurements of the RP release rates as well as the $FL_S$ and RP content for these materials are described below (**Fig. S2, Fig. S3,** and **Table S1**).

3.6 *In vitro* RP release rates from $FL_s$-MS~$RP_R$

Release of RP was measured using a previously reported assay (6). In brief: samples (0.150 mL) of $FL_s$-MS~$RP_R$ (**2A**, **2B**, and **2C**) slurry were placed into nylon mesh pouches and then the pouches were heat sealed. The pouches were placed into divided cuvettes containing 2.7 mL of pH 8.4 bicine buffer (0.1 M) at 37 $^oC$ and a magnetic stir bar. The absorbance of the buffer was measured over 35 h using a heated and stirred auto sampler on a UV-VIS spectrophotometer (Hewlett Packard 8453). RP (565 nm) was released over the course of monitoring, and as expected, no release of the stably linked $FL_S$ (495 nm) was observed. The time to reverse gelation ($t_{RG}$) for the fluorescein labeled gels is 200 h at pH 8.4

(4) and should not occur over the duration of this experiment. Half-lives for RP release were determined using a first order exponential fit of the experimental data (**Fig. S2**). Half-lives measured at pH 8.4 were converted to pH 7.4 values based on $T_{1/2\_7.4} = T_{1/2\_8.4} * 10^{(8.4-7.4)}$ (3).

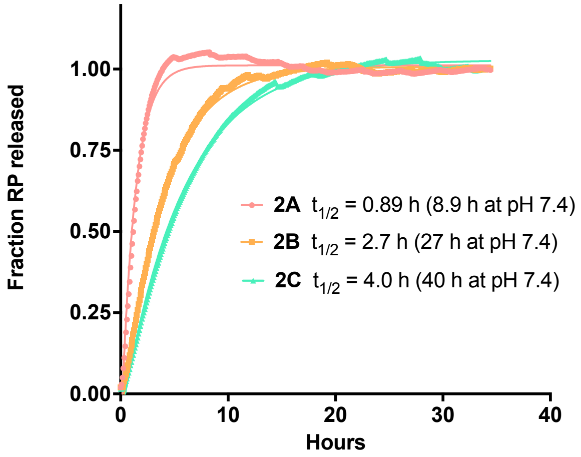


**Figure S2.** Release of RP from $FL_s$-MS~$RP_R$ at pH 8.4, 37 °C showing calculated half-lives for materials **2A**, **2B** and **2C**.

#### 3.7 Dosing syringes, dosing accuracy, and $FL_s$/RP ratio for $FL_s$-MS~$RP_R$

Slurries of $FL_s$-MS~$RP_R$ **2A**, **2B**, and **2C** were passaged once through a 25 gauge tube (Cadence Micro-Emulsification tube, 7974) connected between two 10 mL Luer lock syringes to ensure they were fluid suspensions void of aggregates. Then 0.070 mL of slurry was transferred to a 0.250 mL glass syringe (Hamilton Gastight 1725) using a female-female Luer coupling (Cole Palmer). The glass syringe was fitted with a 30 g x 1/2" needle (BD 305106) then purged by expelling all but 0.025 mL of material through the needle. Four syringes containing 0.025 mL of MSs were prepared for each material (**Fig. S3A**). The needle was capped with a small piece of platinum cure silicone cord (McMaster Carr 9808K21) to prevent leakage and evaporation prior to use. To assess the dosing accuracy the material in the syringes (0.025 mL) was injected into a 1.5 mL centrifuge tube containing 1.0 mL of sodium hydroxide (0.1 N). After 1 hour the absorbance of the solution was measured to detect $FL_s$ (495 nm) and RP (565 nm). The molar ratio of $FL_S$ to RP was determined for each material using $A_{495}$ and $A_{565}$ (**Table S1**). Dosing syringes for use in intra-tumoral injections were prepared in the exact same manner in a laminar flow cabinet, and using syringes that had been disinfecting with 70% ethanol.

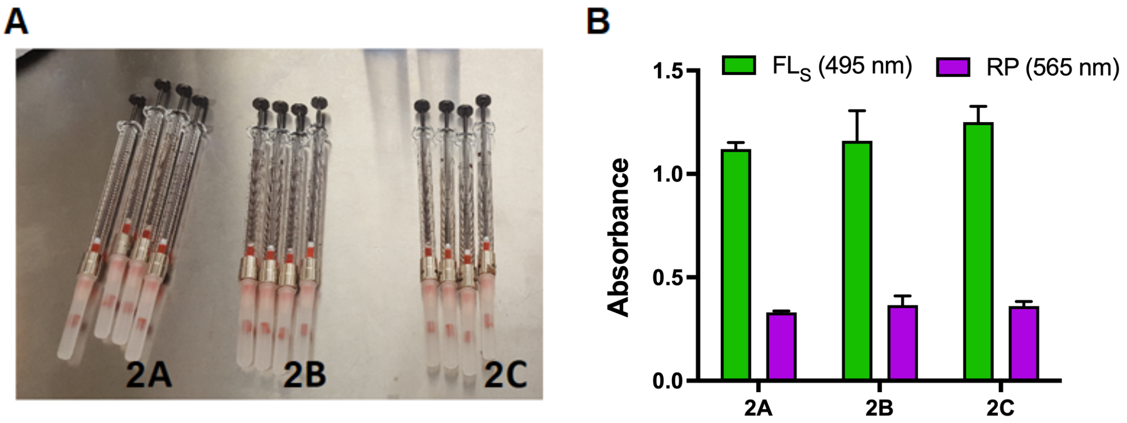

**Figure S3.** Dosing syringes and dosing accuracy. A) Dosing syringes containing 0.025 mL single doses of materials **2A**, **2B** and **2C** with purged and capped needles. B) Average absorbance of $FL_S$ and RP from multiple (N = 4) 0.025 mL doses of **2A**, **2B**, and **2C** into 0.1M NaOH error bars are standard deviation.

**Table S1**. Accuracy of dispensing 0.025 mL of MSs from a 250 μL syringe with a 30 g needle

| Material | FL A495 | SD | CV | | RP A565 | SD | CV | | FL/RP | SD | CV |
|---|---|---|---|---|---|---|---|---|---|---|---|
| RP | NA | NA | | | 0.386 | 0.05 | 0.13 | | --- | --- | --- |
| 2A | 1.125 | 0.03 | 0.03 | | 0.330 | 0.01 | 0.03 | | 4.1 | 0.17 | 0.04 |
| 2B | 1.161 | 0.15 | 0.13 | | 0.365 | 0.05 | 0.14 | | 3.8 | 0.72 | 0.19 |
| 2C | 1.248 | 0.08 | 0.06 | | 0.361 | 0.02 | 0.06 | | 4.2 | 0.35 | 0.08 |

### 4. Assay development, and intra-tumoral injections of $FL_s$-MS~$RP_R$

#### 4.1 *In vitro* separation of RP from $FL_s$-MS-Ac following blending

Mixture of free RP and $FL_S$-MS-Ac: RP (0.122 mL, 5.9 mM, 0.72 nmol) in IAT buffer was mixed into a slurry (3.48 mL) of $FL_s$-MS (0.8 mM $FL_S$) then the material was loaded into insulin syringes (BD 324702) containing 0.100 mL of slurry each.

Separation of RP from $FL_S$-MS-Ac: Two separation methods (centrifugation and filtration) were tested to isolate $FL_s$-MS-Ac and free RP. Both methods began by injecting 0.10 mL of slurry into IAT buffer (1.0 mL), the resulting suspension was homogenized with a tissue blender (IKA T25 Ultra-Turrax homogenizer) for 1 minute at 8000 RPM

Centrifugation Method: Homogenized suspensions were centrifuged 20000 x G for 2 min, then the supernatant extract was removed by pipette. The pellet of $FL_s$-MS-Ac was washed with 5 x 1 mL of IAT, separating MSs by centrifugation at 20000 x G for 2 min between washes.

Filtration method: Homogenized suspensions were filtered using 0.2 μm PTFE spin filters (Millipore Ultrafree MC hydrophilic, Cat no. UFC30LG25) by centrifugation at 10000 x G for 5 min. The filtrate extract was collected and the retained pellet of $FL_s$-MS-Ac was washed with 4 x 0.5 mL of IAT, by spinning at 10000 x G for 5 min for each wash.

Analysis of pellets and washes from both methods (**Fig. S4**): After the final wash the pellet was dissolved in 1.00 mL of 1N NaOH for 1 h. The supernatant (or filtrate) extracts and wash samples were diluted 1:1 with 1N NaOH and kept for 1 h. Next all samples were diluted 1:10 with 1N NaOH then a portion (0.200 mL) was transferred to a well of a black microtiter plate. Fluorescence was measured using a plate reader as described in the general methods.

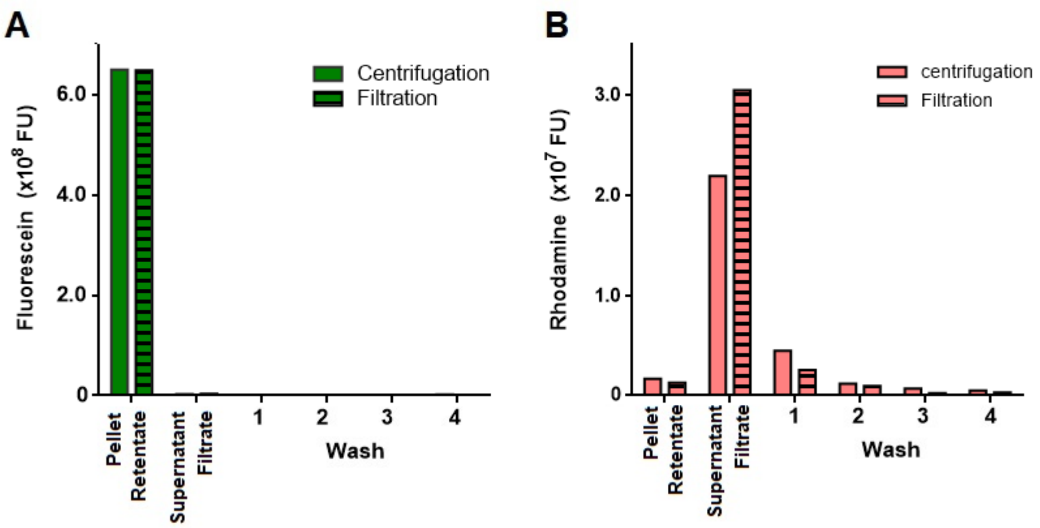


**Figure S4.** Separation of $FL_S$-MSs (A) from free RP (B) after blending with a tissue homogenizer.

#### 4.2 Clearance half-life of RP in tissue, following SC injection into rats

After shaving, sub cutaneous injections of the RP/$FL_S$-MS-Ac mixture in insulin syringes described above (0.100 mL each) were made into the backs of 3 female Sprague Dawley rats (body weight ~250 g) according to the schedule and diagram in **Table S2.** After injection the implants were marked with a permanent marker (Sharpie) by drawing a 12 mm circle around the implant. After injection #5 the animals were euthanized with $CO_2$ then injection #6 was made immediately prior to harvesting tissue. Tissue was harvested to a depth down to the peritoneal membrane, using a 12 mm biopsy punch (Acuderm Acu.Punch P1250). Tissue samples were placed into 2 mL screw cap vials and stored at -80 $^o$C prior to analysis. Untreated tissue samples (X) were also collected to serve as background samples.

**Table S2.** Dosing schedule and injection map for implantation of RP/$FL_S$-MS mixture into rats.

| Implant | Dose time | Hours prior to harvest |
|---|---|---|
| 1 | 10:00 AM | 2.5 |
| 2 | 10:30 AM | 2.0 |
| 3 | 11:00 AM | 1.5 |
| 4 | 11:30 AM | 1.0 |
| 5 | 12:00 PM | 0.5 |
| --- | 12:15 pM (euthanize) | --- |
| 6 | 12:30 PM | 0 |

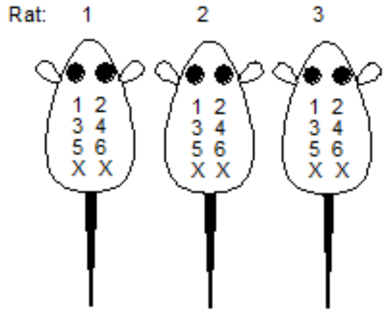


#### 4.3 Determination of $FL_S$ and RP in tissue

Tissue samples were thawed then placed into 5 mL snap-cap round bottom centrifuge tubes. Tissue samples were covered with 2.00 mL of IAT buffer, homogenized at 8000 RPM for 1 minute (IKA T25 Ultra-Turrax homogenizer) then allowed to stand for 20 min prior to centrifugation at 5000 x G for 20 min. The supernatant was transferred to a 5 mL Eppendorf tube. The pellet was treated with an additional 2.00 mL of IAT, mixed, allowed to stand for 20 min, then centrifuged at 5000 x G for 20 min. The supernatant was removed and combine with the first supernatant. The pellet was digested in 4.0 mL of 1 N NaOH for 1 h at room temperature. The pellet digest and supernatant (extract) were diluted to exactly 5.0 mL with water. For $FL_S$ detection samples were further diluted 5 fold with water and for RP detection samples were measured as is. Samples were added to a black 96-well microtiter plate (0.250 mL/well) and analyzed for fluorescence as described in the general methods (**Fig. S5**).

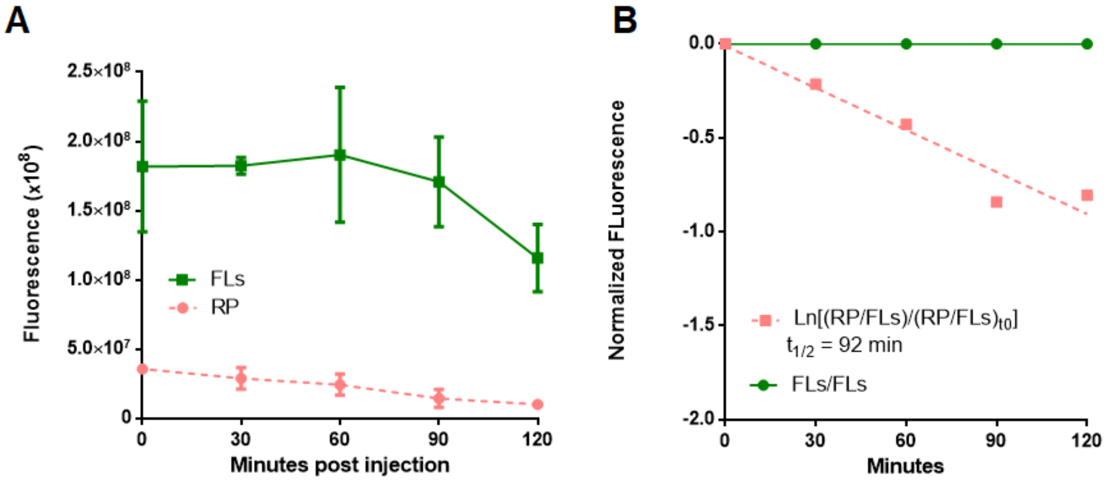


**Figure S5.** Plot of free RP and $FL_s$-MS remaining in injection sites following SC injection of a mixture of RP and $FL_S$-MS-Ac into rats. A) Raw data. B) Normalized data using the ratiometric method.

4.4 Tumor xenografts in mice and intra-tumoral injection of $FL_s$-MS~$RP_R$

Female Balb/c mice weighing 21 ± 1.2 g were implanted with $1x10^5$ CT26 Murine Colon Carcinoma cells, sc, in 0.100 mL of serum free media. One implant per animal was made. Tumors were allowed to grow to 251 ± 90 $mm^3$ (12 days) as determined by measurement with calipers prior injections. Intra-tumoral injection of $FL_s$-MS~RPR slurries (**2A**, **2B** and **2C**) were made from 0.250 mL glass syringes with 30 g needles described above (**Fig. S3**). The 0.025 mL payload of each syringe was injected into a single tumor in two portions (~0.0125 mL each) on either side of the tumor. IAT buffer was injected in the same manner to provide background samples.

Tissue harvesting: At the desired time (hours) after injections, animals were euthanized by cervical dislocation, then the tumors and surrounding tissue were harvested using a 12 mm biopsy punch (Acuderm Acu-Punch P1050). Tissue samples were placed into 10 mL round bottom snap cap centrifuge tubes (Green Bio Research MC01100) containing 4 mL of pH 5.0 IAT buffer. Samples were stored at 4 °C for 40 min prior to being frozen on dry ice then stored at -80 °C for 24 h prior to assay.

4.5 Determination of $FL_S$ and RP in excised tumors

Tissue samples were thawed to room temperature and immediately homogenized at 8000 RPM for 1 minute (IKA T25 Ultra-Turrax homogenizer). Samples were then centrifuged at 5000 x G for 30 min. The supernatant extract (~4 mL) was transferred to a 15 mL Falcon tube. The pellet washed twice IAT buffer by treating with 2.00 mL of buffer, allowed to rock for 45 min, then centrifuging at 5000 x G for 20 min. The wash extracts were removed and combined with the original supernatant extract to give (~8 mL) of total extract. Extract solutions were treated with 1 mL of NaOH (1 N) and diluted to 10 mL with water. The pellets were treated with 1 mL of 1 N NaOH and then diluted to 10 mL with water. Standards were made by injecting 0.025 mL of each material (**2A**, **2B** and **2C**) from dosing syringes into 10 mL of NaOH (0.1 N). After standing for 1 hour the samples were centrifuged at 5000 x G for 20 min, then 0.150 mL of each was loaded into a black 96 well microtiter plate and scanned for fluorescence as described in the general methods.

### 5. Release of SN-38 from subcutaneous microspheres

#### 5.1 Preparation of microspheres with releasable SN-38 (MS~SN-38)

**Scheme S4**. Synthesis of microspheres with releasable SN-38A

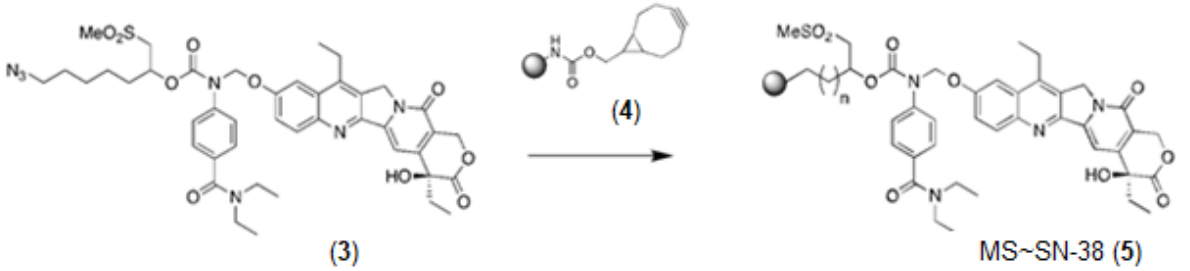


Bicyclononyne-microspheres (BCN-microspheres, **4**) were prepared from 10kDa tetra-PEG amino-MS crosslinked with the $\alpha$-LYS gem dimethyl (GDM) linker containing Mod = $Me_2SO_2N$- (7) using BCN-HSC (Synaffix, SX-A1028). A slurry of amino-MS (53.53 mL, 2.64 mM amine) in 100 mM NaOAc pH 4.0 was placed into a 100 mL GL45 glass bottle and sterilized in and autoclave (5)**.** The sterilized amino-MSs were transferred to a 500 mL conical tube and washed with 2 x 300 mL of sterile aqueous 25% acetonitrile (%v/v), separating the amino-MS from the excess solvent by centrifugation (3700 x G, 10 min) between washes. Prior to washing the acetonitrile was sterilized by filtration using a 0.2 µm PTFE 50 mm Disc filter (Saint Gobain PureFlo, D50CF0201N1N-1), whereas the water was sterilized in the autoclave. The amino-MS were then washed with 5 x 200 mL of acetonitrile as described above to give packed amino-MS slurry in acetonitrile (containing 141 µmol amine). This slurry was then treated with triethylamine (4 eq., 565 µmol) and BCN-HSC (1.2 eq., 170 µmol) in 8 mL acetonitrile. The reaction was mixed end-over-end at ambient temperature for 2 h. at which time a qualitative TNBS test (5) confirmed that the amines had been consumed. Acetic anhydride (1 eq, 141 µmol) was then added for 0.5 h to cap any residual free amines, and the slurry was washed with 6 x 250 mL acetonitrile as described above. The final packed slurry of BCN-microspheres was 50 mL and contained 141 µmol BCN and was used as is for loading with $N_3$-linker($SO_2Me$)-SN-38 (below).

$N_3$-linker($SO_2Me$)-SN-38 (**3**) was synthesized as previously described (8). A 17.3 mM solution of $N_3$-linker($SO_2Me$)-SN-38 (8.95 mL, 155 µmol, 1.1 equiv) in acetonitrile was added to 53.5 g BCN loaded microspheres in acetonitrile (141 µmol BCN, 1.0 equiv). The reaction mixture was mixed by rocking end over end at 37°C for 22 hr. Progress was monitored by following loss of SN38 in the supernatant by $A_{363}$. The loaded microspheres were washed with 4 x 250 mL of acetonitrile followed by 6 x 250 mL of IAT-Met buffer (10 mM NaAcOH, 143 mM NaCl, 0.05% Tween 20, pH 5.0, 10 mM methionine), separating MS from excess buffer by centrifugation (3700 x G, 10min) between washes. The resulting slurry of MS~SN-$38_R$ (**5**) was pelleted (3700 x G, 10 min), transferred to a 10 mL syringe and stored at 4°C. The mass of the packed slurry was 10 g.

The concentration and loading efficiency of the MS~SN-$38_R$ was determined by dissolving 20 µL of the packed slurry (20 mg) in 50 mM NaOH (80 µL) for 1 hour at room temperature. The SN-38 content was determined by absorbance of the SN-38 anion by $A_{414}$. The PEG content in the solution was determined using a previously described colorimetric PEG assay (5). The percent loading of the microsphere slurry was determined to be 98%, by the ratio of SN-38 to PEG (found 196 +/- X nmol SN-38 / mg PEG, 200 theory).

#### 5.2 In vitro release and dissolution of MS~SN-38

Samples of MS~SN-38 (60 µL) were placed in custom made dissolution cells with permeable nylon mesh membranes (**Fig. S6**). These cells are simply smaller scale versions of previously described dissolution cells (5). The cells were place in 5 mL tubes containing 4.1 mL of 100 mM Na borate buffer pH 9.4 pre-equilibrated to 37°C in a temperature-controlled stirred heat block (**Fig. S6**). A liquid handling robot (5) was programmed to remove 60 µL samples from the reaction buffer at t=0 and specified time points over 50 h. The concentration of SN-38 in the reaction supernatant was determined by absorbance at 414 nm based on previously described methods (5) The $A_{414}$ of the supernatant vs time were fit to a single exponential using Prism software to determine the release $t_{1/2}$ for SN-38 was 1.6 h. Similarly, the PEG content of the supernatant was determined using a previously described PEG assay (5) to determine the $t_{rg}$ = 25 h (**Fig. S7**).

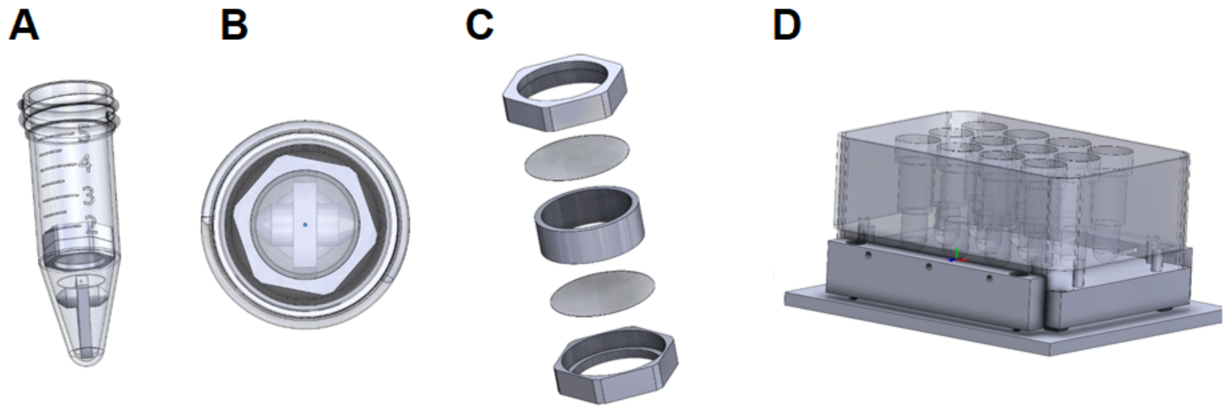


**Figure S6.** Small scale PEEK dissolution cell. A) Cell in 5 mL conical tube with stirrer. (**B**) Top view of the assembly shown in part **A**. C) Exploded view of cell showing body, caps and nylon mesh membranes. D) 12 position stirring heat block for operation of cells on a liquid handling robot.

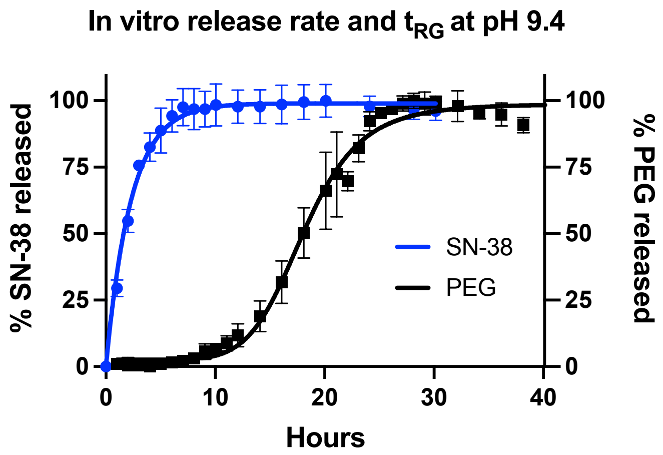


**Figure S7.** *In vitro* release and dissolution of MS~SN-38 at pH 9.4, 37°C. The release $t_{1/2}$ for SN-38 was measured to be 1.6 h at pH 9.4 or 160 h at pH 7.4 based on $T_{1/2\_7.4} = T_{1/2\_9.4} * 10^{(9.4-7.4)}$ (3) and MS dissolution $t_{RG}$ was 25 h at pH 9.4.

### 5.3 *In vivo* release of SN-38 from microspheres

#### 5.3.1 Dose formulation preparation and analysis

MS~SN-38 and $FL_s$-MS-Ac described above were combined to generate a slurry containing 100 μM SN-38 and 32 μM fluorescein in IAT buffer containing 1% 40kDa hyaluronic acid (5). The SN-38 and $FL_s$ concentration of the mixture was determined by dissolving 25 uL samples of the slurry in 0.1 M NaOH (175 μL) for 1 h at room temperature. Fluorescein and SN-38 were quantitated spectrophotometrically at by $A_{495}$ for $FL_s$ and $A_{363}$ for SN-38. This material was loaded into insulin syringes (BD 324702) containing 0.100 mL of slurry each to administer SC injections.

#### 5.3.2 Dose administration and recovery

On Day 0, seven male SD rats were weighed, anesthetized with isoflurane, and their backs shaved. Then, 100 μL of microsphere slurry (containing the mixture of MS~Sn-38 and FLs-MS-Ac described above) was injected SC into 4 sites equally spaced on their backs. The location of each injection was marked with circular tattoo ~12 mm in diameter prior to injection. On days 0, 1, 3, 6, 12, 18 and 24, one rat/day was euthanized with $CO_2$, then the hair was removed by shaving and treatment with Nair (Church & Dwight), followed by cleaning with 70% isopropyl alcohol. Using the tattoos for reference, the tissue surrounding the SC injection site was excised down to the peritoneal membrane using a 12 mm diameter biopsy punch (Acuderm Acu-Punch P1250). For each rat, a control skin sample (no injection) was obtained from the flank at least 2 cm away from all injection sites. Each tissue sample was placed into a 1.5 mL Eppendorf tube and stored at -80 °C until analysis.

#### 5.3.3 Analysis of free SN-38 and MS-bound SN-38 at injection sites

To assess free SN-38 present at the injection site tissue samples were thawed in 1 mL 0.5% HOAc, then mixed with a Dounce homogenizer. The pestle was rinsed with 1 mL of 0.5% HOAc which was combined with the homogenized tissue sample. Samples were then clarified by centrifugation (15 min) and the supernatant was removed. The tissue sample was washed with

2 mL 0.5% HOAc, clarified by centrifugation (15 min) and the wash was combined with the previous supernatant to give ~4 mL 0.5% HOAc extract for analysis. A sample of each extract (0.050 mL) was analyzed by HPLC (below) to determine free SN-38 present in the tissue surrounding the injection site.

To assess the content of MS~SN-38 and MS-$FL_s$-Ac at the injection site, the tissue pellets from the HOAc extracts (above) were treated with 0.25 M NaOH (4 mL) vortexed to mix then heated to 80 °C for 1 h to dissolve MS and release MS bound SN-38. The mixture was clarified by centrifugation (15 min), and the supernatant was removed for analysis. A sample of each supernatant (0.050 mL) was analyzed by HPLC to quantify SN-38 and $FL_s$.

HPLC analysis. The acid and base extracts of tissue samples described above were analyzed by reverse phase HPLC using the following method: 0% ACN 0-1min, 0-100% 1-30 min, 100% 30-31min, 100-0, 31-32min, 0% 32-33 min all with 0.1% TFA. Fluorescence detection: EX 360nm EM 545nm (0-14.5 min) for SN-38, and EX 442nm EM 520nm, (14.5-30 min) for Fluorescein. SN-38 eluted at 13.2 min, and PEG-fluorescein eluted at 16.7 min. An SN-38 standard curve was generated using 50 uL 0.72 nM to 11.3 uM SN-38 quantitated by HPLC. The fluorescent SN-38 peak area of 50 µL injections was plotted vs. concentration to obtain the standard curve.

#### 5.4 Anti-tumor efficacy of MS~SN-38

The 22Rv1 ATM KO cell line (9) was tested negative for *Mycoplasma* contamination prior to the initiation of in vivo studies. Tumor xenograft studies were conducted at Murigenics. Male NSG mice were injected SC with 22Rv1 ATM KO cells (1 x$10^7$ in 100 µL of 1:1 PBS:Matrigel). When the average tumor volume reached ~125 $mm^3$, MS~SN-38 (0.2 – 7.5 µmol/kg) was administered IT (50 µL injection) or SC (50 µL injection) and PLX038A (7.5 µmol/kg) was administered IP (**Fig. 4, S8**). The control group was left untreated until the average tumor volume reached ~1,000 $mm^3$. Then, mice were treated with a single IT dose of MS~SN-38 (2 or 20 µmol/kg, 100 µL injection). In combination experiments with TLZ, mice were treated with IT MS~SN-38 (0.2 µmol/kg), SC MS~SN-38 (2 µmol/kg), PO TLZ (0.4 µmol/kg) QDx21 or a combination of IT MS~SN-38 and PO TLZ. The tumor volume (caliper measurement, V= 0.5x(length x $width^2$) and body weights were measured twice weekly.

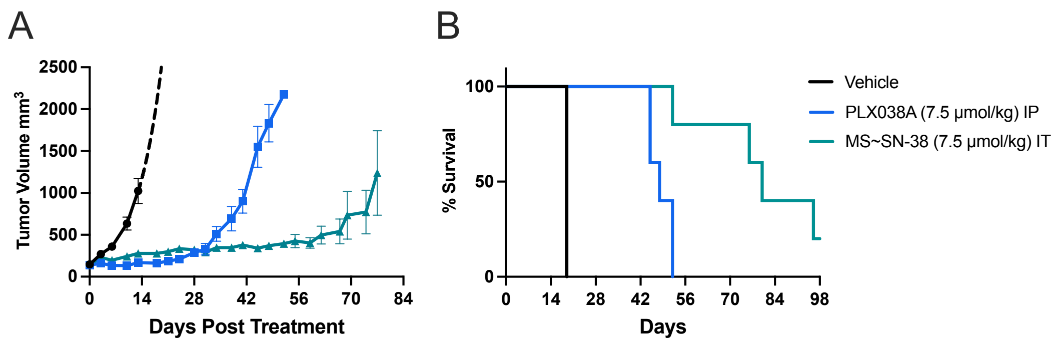


**Figure S8.** Anti-tumor effects of MS~SN-38 and PLX038A in mice bearing 22Rv1 $ATM^{(-/-)}$ xenografts. A) Tumor volume vs days post treatment; control (●) with estimated exponential tumor growth after 1,000 $mm^3$ (dashed line); IT 7.5 µmol/kg MS~SN-38 (▲), and IP 7.5 µmol/kg PLX038A (■). B) Kaplan Meier plot of overall survival for animals treated in A. Tumor volume data represent median tumor volume ± interquartile range (n=5/group).

5.5 Animal Welfare Statement

All animal handling and care was performed by MuriGenics (Vallejo, CA). All experiments were performed using protocol and conditions that conformed to their Institutional Animal Care and Use Committee recommendations.
